# High antibody titres induced by protein subunit vaccines against Buruli ulcer using *Mycobacterium ulcerans* antigens Hsp18 and MUL_3720

**DOI:** 10.1101/2020.02.16.951533

**Authors:** Kirstie M. Mangas, Nicholas Tobias, Estelle Marion, Jérémie Babonneau, Laurent Marsollier, Jessica L. Porter, Sacha J. Pidot, Chinn Yi Wong, David C. Jackson, Brendon Y. Chua, Timothy P. Stinear

## Abstract

**Background:** *Mycobacterium ulcerans* is the causative agent of a debilitating skin and soft tissue infection known as Buruli ulcer (BU). There is no vaccine against BU. The purpose of this study was to investigate the vaccine potential of two previously described immunogenic *M. ulcerans* proteins, MUL_3720 and Hsp18, using a mouse tail infection model of BU.

**Methods:** Recombinant versions of the two proteins were each electrostatically coupled with a previously described lipopeptide adjuvant. Seven C57BL/6 and seven BALB/c mice were vaccinated and boosted with each of the formulations. Vaccinated mice were then challenged with *M. ulcerans* via subcutaneous tail inoculation. Vaccine performance was assessed by time-to-ulceration compared to unvaccinated mice.

**Results:** The MUL_3720 and Hsp18 vaccines induced high titres of antigen-specific antibodies that were predominately subtype IgG_1_. However, all mice developed ulcers by day-40 post-*M. ulcerans* challenge. No significant difference was observed in the time-to-onset of ulceration between the experimental vaccine groups and unvaccinated animals.

**Conclusions:** These data align with previous vaccine experiments using Hsp18 and MUL_3720 that indicated these proteins may not be appropriate vaccine antigens. This work highlights the need to explore alternative vaccine targets and different approaches to understand the role antibodies might play in controlling BU.

## Introduction

Buruli ulcer (BU) is a disease caused by *Mycobacterium ulcerans. M. ulcerans* infects subcutaneous tissue and commonly presents as a skin nodule (in Africa) or papule (in Australia), sometimes accompanied by redness; however, oedema is another common initial presentation. As the disease progresses the skin around the infected area breaks down and an ulcer develops [1, 2]. Ulcers typically present with deep undermined edges and have a necrotic core comprised of slough of bacteria, dead skin and immune cells [3, 4]. Infections are rarely fatal but untreated ulcers can destroy fat tissue, blood vessels, muscles and bone [5, 6].

The transmission of BU is likely caused by the introduction of *M. ulcerans* beneath the skin. This could be achieved through the puncture of *M. ulcerans-*contaminated skin (with examples in the literature of infections following human bite, bullet and land mine wounds, or vaccination) or by the introduction of *M. ulcerans* contaminated objects into the subcutaneous tissue, such as following insect bites [7-9]. BU endemic areas are focused in certain rural regions across west, sub-Saharan and central Africa, including Nigeria, Ghana, Togo, Cameroon, Benin, Democratic Republic of Congo and Côte d’Ivoire. The disease also occurs in Australia – primarily on the Bellarine and Mornington Peninsulas near the major metropolitan centre of Melbourne [10-12]. The disease can affect all age groups and ethnicities [13]. In Australia, ulcers are predominately reported on upper (27%) and lower limbs (70%) [14].

*M. ulcerans* is a slow-growing bacterium, with a doubling time of greater than 48 hours. As such, symptoms of BU can take months to appear after primary infection. If diagnosed early, BU can be treated effectively by combination antibiotic therapy [15]. Unfortunately, in many cases the disease can initially be misdiagnosed as other more common skin infections [16, 17]. Delayed diagnosis and treatment can lead to extensive lesions that leave victims with life-long disfigurement and disability. Reparative surgery is often required for severe cases [18]. A retrospective study in Australia showed that most diagnoses (87%) occurred once ulceration has been reached [19] and in Ghana 66% cases were diagnosed with active lesions [20]. There is currently no protective treatment for BU and no distinct mechanism of transmission. Furthermore, treatment can be difficult to access for those in rural areas. Thus, there is a need to develop an effective vaccine to protect those particularly in highly endemic areas.

The *M. bovis* ‘BCG’ vaccine has been shown to delay the onset of BU symptoms and decrease bacterial load in both experimental animal BU infection models and in studies of human populations [21-25]. Therefore, the BCG vaccine is the benchmark for assessing potential *M. ulcerans* vaccines. Some studies have assessed the efficacy of putative BU vaccines although none have reached clinical trials [21, 22, 26-37]. All these vaccines were tested in murine challenge models and were not capable of preventing the eventual onset of disease.

One approach to vaccination is to use antigens specific for a specific pathogen (e.g. certain proteins(s) that are recognized by the immune system and induce neutralizing antibodies [38, 39]. For rapid immune recognition these proteins would ideally be cell surface associated. Two *M. ulcerans* proteins MUL_3720 and Hsp18 have been identified as potential candidates for vaccine antigens. Hsp18 is a protein associated with biofilm formation and *M. ulcerans*-infected individuals produce antibodies against Hsp18 [40, 41]. MUL_3720 is a highly expressed cell-wall associated protein with a putative role in cell-wall biosynthesis [42, 43].

As protein antigens may be poorly immunogenic on their own, adjuvants are used to enhance antigenic potency. A lipopeptide adjuvant known as R_4_Pam_2_Cys has been found to increase antigen uptake, increase dendritic cell trafficking to lymph nodes and enhance antibody production against antigens derived from pathogens including influenza and hepatitis C in murine models [44-47].

The aim of this study was to try to develop a preventative vaccine against Buruli ulcer, comprising two highly expressed cell-wall associated proteins, MUL_3720 or Hsp18, bound to an R_4_Pam_2_Cys-based lipopeptide adjuvant.

## Materials & Methods

### Strains and culture conditions

*Escherichia coli* Rosetta2 containing plasmid pET30b-Hsp18 (strain TPS681) or pDest17-MUL_3720 (strain TPS682) was grown at 37°C in Luria-Bertani (LB) broth (Difco, Becton Dickinson, MD, USA) supplemented with 100 µg/ml ampicillin (Sigma-Aldrich, USA) or 50 µg/ml kanamycin to express 6xHIS-tagged Hsp18 or MUL_3720 recombinant protein. *Mycobacterium ulcerans* (strain Mu_1G897) was grown at 30°C in 7H9 broth or 7H10 agar (Middlebrook, Becton Dickinson, MD, USA) supplemented with oleic acid, albumin, dextrose and catalase growth supplement (OADC) (Middlebrook, Becton Dickinson, MD, USA), and 0.5% glycerol (v/v). *M. bovis* BCG (strain Sanofi Pasteur) used for vaccinations was grown at 37°C in 7H9 broth or 7H10 agar supplemented with OADC. Mycobacterial colony counts from cultures or tissue specimens were performed using spot plating as previously described [48].

### Recombinant protein expression

Overnight cultures of strains TPS681and TPS682 were diluted to OD_600_ = 0.05 in LB broth. Each culture was incubated at 37°C with shaking at 200 rpm until OD_600_ = 0.6-0.7, then 1 mM IPTG (Isopropyl b-D-1-thiogalactopyr-anoside) was added to induce protein expression. The cells were incubated for a further four hours to express the protein. To harvest the protein, cells were resuspended in wash buffer (8 M urea, 150 mM sodium chloride, 10% glycerol) and sonicated at amplitude 60 (QSonica Ultrasonic Liquid Processor S-4000, Misonix) until the solution turned clear. The lysate was filtered with a 0.22 µM filter (Millipore) to remove cellular debris and the protein was column-purified using anti-histidine resin (ClonTech). The resin was washed ten times with 10x column volumes of wash buffer mixed with an increasing proportion of tris buffer (20 mM Tris-HCl, 150 mM sodium chloride, 10% glycerol) until the column was washed with only tris buffer. The resin was washed a further two times with tris buffer containing 20 mM imidazole. Protein was eluted in tris buffer containing 200 mM imidazole and dialysed in phosphate buffered saline (PBS) before concentration using a 3K MWCO PES concentration column (Pierce).

### Sodium dodecyl sulphate polyacrylamide gel electrophoresis (SDS-PAGE)

Samples were denatured in an equal volume of 2 x sample loading buffer (40% (v/v) 0.5M Tris-HCL pH 6.8, 10% glycerol, 1.7% (w/v) SDS, 10% 2-β-mercaptoethanol, 0.13% (w/v) bromophenol blue in distilled water) at 100°C for 5 minutes. Ten microlitres of each sample and SeeBlue® Plus2 pre-stained protein standard (Invitrogen) were loaded into a 0.5mm 12% polyacrylamide gel under reducing conditions, as previously described [49]. The gel was run in running buffer (0.3% (w/v) Tris, 1.44% (w/v) glycine and 0.1% (w/v) SDS in distilled water) for 1 hour at 150 volts (Mini-protean vertical electrophoresis cell, Bio-Rad). The gels were stained in Coomassie stain (45% methanol, 10% acetic acid 0.25% (w/v) Coomassie brilliant blue in distilled water) for 1 hour and destained in Coomassie destain (33% Methanol, 10% acetic acid, 60% distilled water) until the protein bands could be identified.

### Western Blotting

Proteins were separated on a 12% polyacrylamide gel as per the method for SDS-PAGE. After separation proteins were transferred to a nitrocellulose membrane in tris-glycine transfer buffer (1.5 mM Tris, 12mM glycine, 15 % methanol (v/v) in distilled water) for 1 hour at 100 volts (Mini Trans-Blot Cell, Bio-Rad). The nitrocellulose membrane was blocked in blocking buffer (5% (w/v) skim milk powder and 0.1% Tween-20 in PBS) overnight at 4°C. The membrane was incubated in blocking buffer containing anti-6xHIS-HRP antibody (Roche Applied Science) at 1:500 dilution. The membrane was washed in PBS containing 0.1% Tween-20 and then exposed to developing solution (Western Lighting Chemiluminescence kit, Perkin Elmer) according to manufacturer’s guidelines. Chemiluminescence was detected using an MF ChemiBIS gel imaging system (DNR Bio-Imaging Systems).

### Analysis of electrostatic interaction between protein antigen and lipopeptide formulations

The association between each protein and R_4_Pam_2_Cys was measured by mixing 25 µg of protein with increasing amounts of lipopeptide in 50 µl PBS in a 96-well plate (Nunc, Thermo Scientific). The formation of protein-lipopeptide complexes through electrostatic interaction was measured by an increase in light absorbance. Plates were read at dual wavelengths of 505 and 595 nm on plate reader (LabSystems Multiskan Multisoft microplate reader).

### Lipopeptide vaccine preparation

Each vaccine dose contained 25 µg protein added to R_4_Pam_2_Cys at a ratio of 1:5 mole of protein to lipopeptide. PBS was added to a final volume of 100 µl and the combination sonicated in a water bath for 30 seconds. Control vaccine preparations were made containing 25 µg protein alone or R_4_Pam_2_Cys lipopeptide alone and sonicated before administration.

### Ethics statement for animal experiments

All animal experiments were performed in full compliance with national guidelines (articles R214-87 to R214-90 from French “rural code”) and European guidelines (directive 2010/63/EU of the European Parliament and of the council of September 22, 2010 on the protection of animals used for scientific purposes). All protocols were approved by the Ethics Committee of region Pays de la Loire under protocol nos. CEEA 2009.14 and CEEA 2012.145. Animals were maintained under specific pathogen-free conditions in the animal house facility of the Centre Hospitalier Universitaire, Angers, France (agreement A 49 007 002). Six-week old female C57BL/6 and BALB/c mice were obtained from Charles River Laboratories (Saint-Germain-Nuelles, France) and housed at CHU Angers. Food and water were given *ad libitum*.

### Vaccination of animals

The synthesis and purification of the branched cationic lipopeptide, R_4_Pam_2_Cys, was performed as previously described [45, 50, 51]. Each vaccine dose contained 25 µg protein formulated in PBS with R_4_Pam_2_Cys at a 1:5 molar ratio of protein to lipopeptide in a final volume of 100 µl. The protein alone control formulation contained 25 ug protein per dose diluted in PBS. The R_4_Pam_2_Cys alone formulations contained the same amount of lipopeptide used in each of the protein + adjuvant formulations, calculated by the 1:5 molecular ratio (with the omission of the protein from the solution). The R_4_Pam_2_Cys alone formulations were diluted to the correct concentration in PBS. Live-attenuated *M. bovis* BCG strain ‘Sanofi Pasteur’ was grown to log phase and stored at −80°C in 20% glycerol until use. Bacteria were washed with PBS and resuspended in 200ul, before administration at 4.7 × 10^5^ bacteria per dose. All vaccines and control formulations were sonicated for 5 minutes in a waterbath sonicator before being administered.

For vaccination using R_4_Pam_2_Cys, animals were inoculated subcutaneously at the base of tail (100µl per dose at 50 µl per flank) and boosted 21 days later with the same formulations. Mice vaccinated with approximately 1 × 10^3^ CFU *M. bovis* BCG resuspended in PBS at the base of tail (100 µl per dose at 50µl per flank).

### M. ulcerans challenge

Mice were challenged on day 35 by subcutaneous injection on the tail with 1 × 10^4^ CFU *M. ulcerans* (Mu_1G897) resuspended in 50 µl PBS. Mice were allowed to recover and monitored for up to 40 days after infection and euthanised when tail ulceration was observed wherein sera were obtained for immunological analysis.

### Serum antibody titre measurements

Serum was prepared from blood obtained from mice at day 0, day 18, day 33 and day 63. Antibody titres were measured using enzyme linked immunosorbent assay (ELISA) as per methods described in [45]. Briefly, ELISA plates (Nunc, Thermo Scientific) were coated overnight with 5 µg protein diluted in PBSN_3_ and blocked with BSA_10_PBS for 2 hours at room temperature. Plates were washed with PBS containing 0.05% Tween-20 (PBST). Neat sera were sequentially diluted in BSA_5_PBST and incubated at room temperature for 6 hours. Bound antibody was detected by adding horse radish peroxidase conjugated rabbit anti-mouse IgG (Dako, Glostrup, Denmark) at a concentration of 1:400 in BSA_5_PBST for 2 hours. Plates were developed with developing solution (hydrogen peroxide, citric acid and ABTS) and incubated for 10-15 min with gentle agitation to observe a colour change. The reaction was stopped with 50 mM sodium fluoride. Plates were read at dual wavelengths of 505 and 595 nm on plate reader (LabSystems Multiskan Multisoft microplate reader).

### Statistical analysis

Graphpad Prism software (GraphPad Software v7, CA, USA) was used to perform statistical analyses on the antibody titre. Antibody titres were analysed using two-way ANOVA with Tukey’s correction for multiple comparisons. The time to ulceration data were displayed as a Kaplan-Meier plot and statistical significance was determined using a Log-Rank (Mantel-Cox) test. For all tests \**p* < 0.05, \*\**p* < 0.01 and \*\*\**p* < 0.001 and **** *p* < 0.0001 were considered statistically significant.

## Results

MUL_3720 and Hsp18 have previously been shown to be immunogenic and cell-wall associated [40, 43]. The adjuvant Pam_2_Cys has been shown to induce strong antibody responses to proteins from infectious agents such as influenza and hepatitis C [52-54]. Therefore, this study measures the ability of MUL_3720 and Hsp18 based vaccines, incorporating the adjuvant Pam2Cys, to generate protein-specific antibodies and to protect against BU.

### Recombinant MUL_3720 and Hsp18 both bound to R_4_Pam_2_Cys

Recombinant MUL_3720 and Hsp18, expressed from inducible *E. coli* expression vectors, were prepared for use as antigens in the vaccine formulations (Table S1). Purification of the recombinant proteins was confirmed by SDS-PAGE and Western blot analyses of the eluate (Fig. 1). DLS analysis was then performed to identify whether recombinant MUL_3720 or Hsp18 would electrostatically bind to either the positively charged lipopeptide adjuvant R_4_Pam_2_Cys, or its negatively charged counterpart, E_8_Pam_2_Cys. The optical density of solutions containing these constituents at a wavelength of 450nm (OD_450_) is related to the particle size of molecules in solution, reflecting the strength of the ionic interaction between protein and lipopeptide [45]. MUL_3720 preferentially bound to R_4_Pam_2_Cys compared to E_8_Pam_2_Cys (Fig. 2A). This is shown as a gradual increase in optical density following the addition of increasing amounts of R_4_Pam_2_Cys to a constant amount of MUL_3720. At a 5-fold molar excess of protein to lipopeptide the OD_450_ plateaued, suggesting MUL_3720 bound most strongly to R_4_Pam_2_Cys at a 1:5 protein to lipopeptide ratio. Conversely, when E_8_Pam_2_Cys was added to MUL_3720 the optical density remained static and did not increase with increasing lipopeptide concentrations, indicating a lack of binding. Hsp18 also appeared to bind preferentially to R_4_Pam_2_Cys and also at a 1:5 ratio of Hsp18 to R_4_Pam_2_Cys (Figure 2B). Therefore, two protein-adjuvant formulations were prepared using MUL_3720 with R_4_Pam_2_Cys and Hsp18 with R_4_Pam_2_Cys, both at a 1:5 protein to lipopeptide molar ratio.

**Figure 1.**
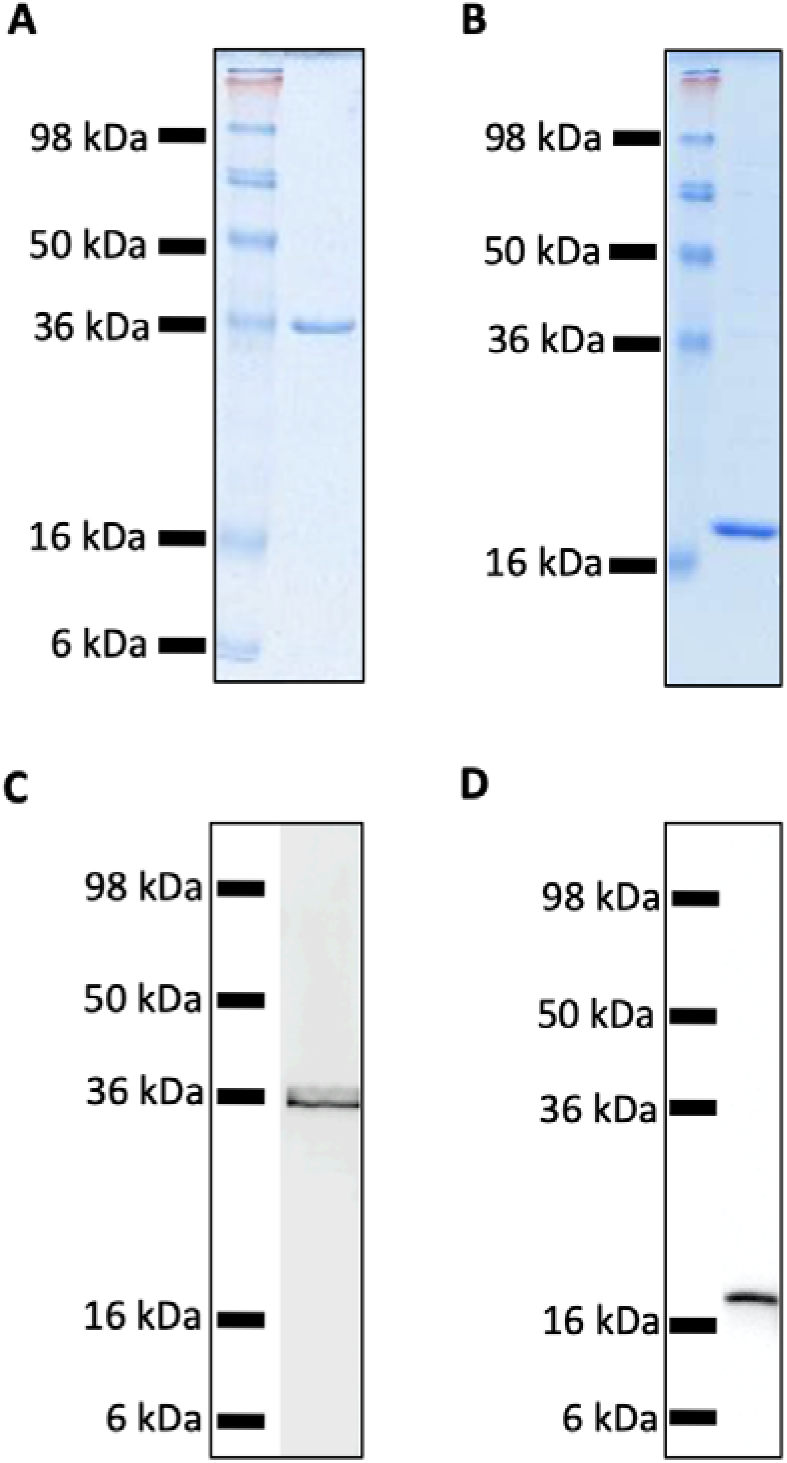
SDS-PAGE and Western Blot Analysis of purified recombinant MUL_3720 and Hsp18 proteins. **(A)** SDS-PAGE of MUL_3720 protein elution (containing 10 µg protein) shows a band ∼36 kDa. (**B)** SDS-PAGE of Hsp18 protein elution (containing 10 µg protein) shows a band ∼18 kDa. **(C)** Protein in the final MUL_3720 elute was analysed by Western Blot using an anti-6xHIS-tag antibody to detect the presence of a single band corresponding to the band as the SDS- PAGE analysis. (**D)** Protein in the final Hsp18 elute was analysed by Western Blot using an anti-6xHIS-tag antibody to detect the presence of a single band corresponding to the 18 kDa band as the SDS-PAGE analysis.

**Figure 2.**
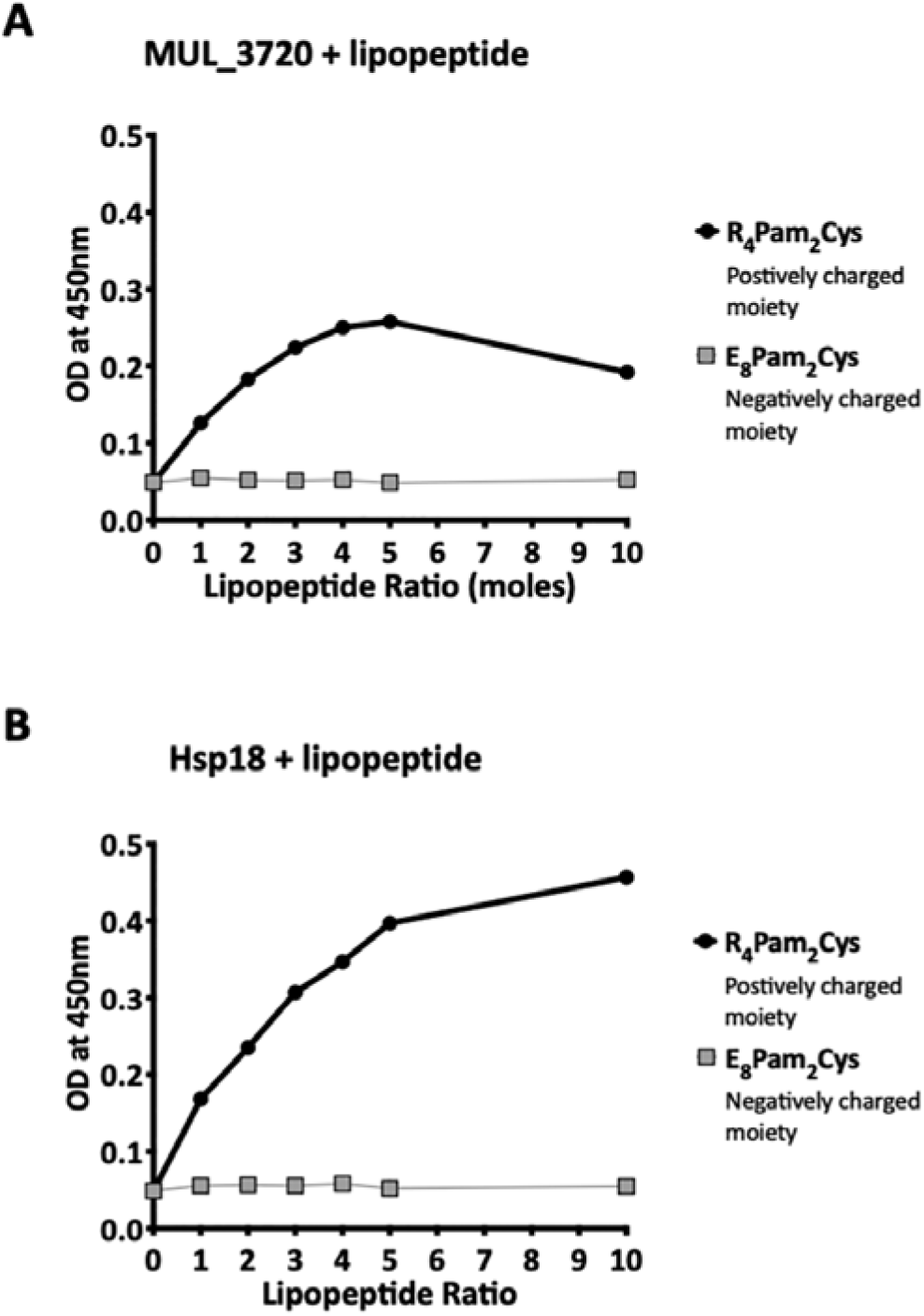
Recombinant MUL_3720 and Hsp18 protein formulation linked with R_4_Pam_2_Cys. To analyse the formation of antigen-lipopeptide complexes, a constant amount of antigen **(A)** MUL_3720 (25µg) and **(B)** Hsp18 (25µg) was mixed with lipopeptide at different protein:lipopeptide molar ratios in 50 µl of PBS. These graphs depict the absorbance values of these solutions at an optical density of 450nm (OD_450_). In these assays either R_4_Pam_2_Cys or E_8_Pam_2_Cys lipopeptides were added to the proteins at increasing amounts. The addition of R_4_Pam_2_Cys is depicted with black circles and the addition of E_8_Pam_2_Cys is depicted with grey squares. An increase in absorbance in correlation to an increase in lipopeptide was indicative of protein binding to lipopeptide.

### Vaccination induced strong protein-specific antibody responses

Prior to challenge with *M. ulcerans*, the ability of the vaccine candidates to generate murine immune responses was assessed. ELISAs were utilized to measure the antibody (IgG) titres in sera obtained from two strains of mice (BALB/c and C57BL/6) immunized with either MUL_3720 + R_4_Pam_2_Cys or Hsp18 + R_4_Pam_2_Cys after the primary vaccination dose (day 18) and a secondary dose (day 33).

Vaccination with MUL_3720 recombinant protein alone or MUL_3720 + R_4_Pam_2_Cys were capable of inducing MUL_3720-specific antibody titres in both BALB/c and C57BL/6 strains of mice (Fig. 3A, B). Primary vaccination with MUL_3720 protein alone induced MUL_3720-specific antibody responses that significantly increased (*p* < 0.0001) following a vaccine boost (*p* = 0.0234). Additionally, MUL_3720 + R_4_Pam_2_Cys generated MUL_3720 specific antibody responses after primary vaccination (*p* < 0.0001 in BALB/c and C57BL/6), which were increased after the secondary boost (*p* < 0.0001 in BALB/c and not statistically significant in C57BL/6). The titres after the boost in particular were greater than MUL_3720 alone vaccination (*p* = 0.0031 in BABL/c and *p* = 0.006 in C57BL/6). Mice that were not vaccinated with recombinant MUL_3720 (R_4_Pam_2_Cys alone and BCG) did not have an increase in MUL_3720-specific antibodies compared to naïve mice.

**Figure 3.**
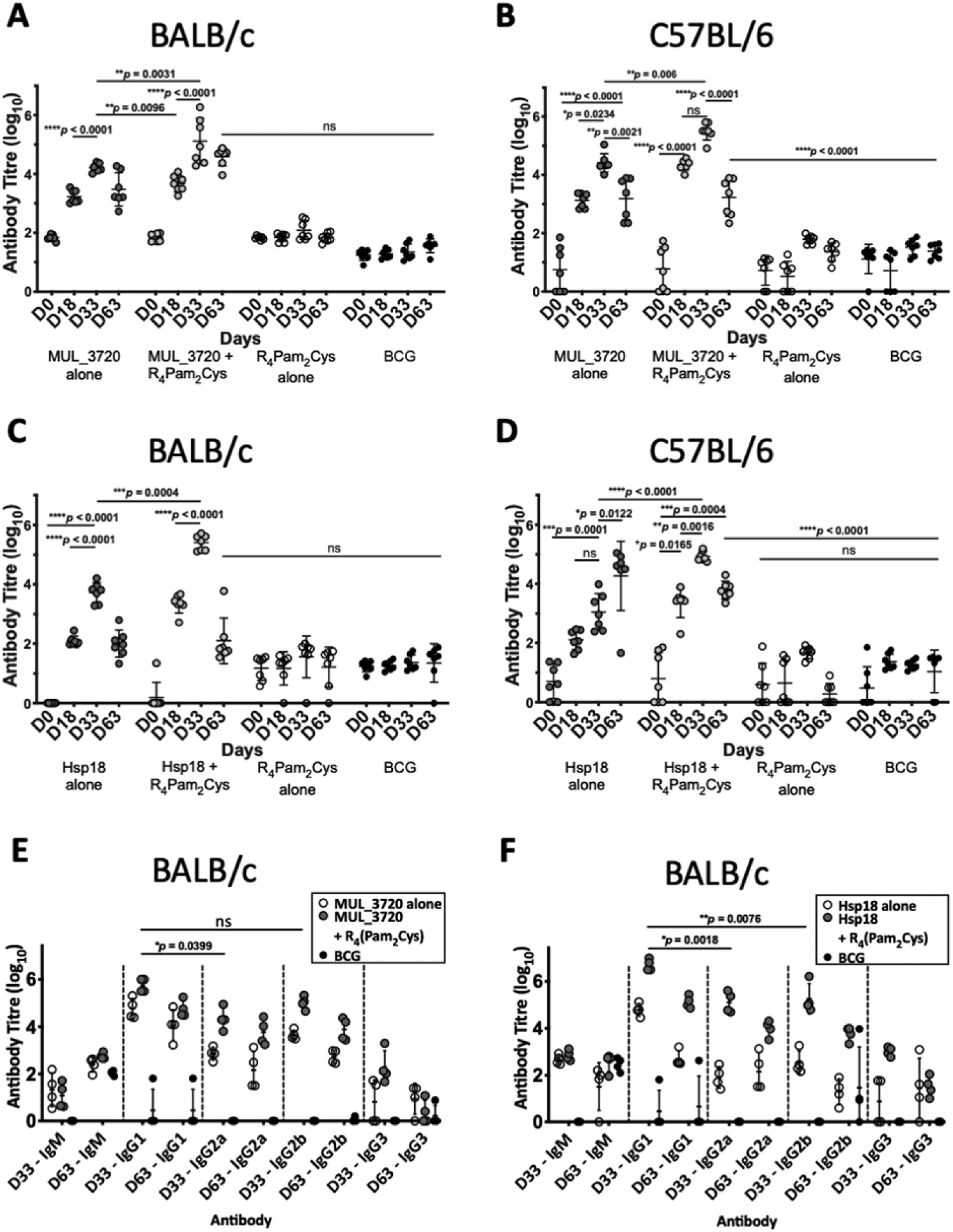
Antibody titres from BALB/c and C57BL/6 mice immunized with recombinant MUL_3720 or Hsp18 linked to R_4_Pam_2_Cys lipopeptide adjuvant. MUL_3720-specific antibody titres from **(A)** BALB/c and **(B)** C57BL/6 mice. Mice were vaccinated with protein alone (MUL_3720) (grey circles), recombinant protein + R_4_Pam_2_Cys (blue circles), R_4_Pam_2_Cys alone (clear circles) and *M. bovis* BCG (black circles). A separate ELISA was performed to measure Hsp18-specific antibody titres in **(C)** BALB/c and **(D)** C57BL/6 mice. Mice were vaccinated with protein alone (Hsp18) (grey circles), recombinant protein + R_4_Pam_2_Cys (blue circles), R_4_Pam_2_Cys alone (clear circles) and *M. bovis* BCG (black circles). IgG isotypes (IgG_1_, IgG_2a_, IgG_2b_ and IgG_3_) were quantified from BALB/c mice immunized with **(E)** MUL_3720 + R_4_Pam_2_Cys and **(F)** Hsp18 + R_4_Pam_2_Cys. Mice were vaccinated with protein antigen alone (either MUL_3720 or Hsp18) (clear circles), protein + R_4_Pam_2_Cys (grey circles) and BCG (black circles). Results are shown as zero if below detectable limits. The null hypothesis (no difference in mean antibody responses between treatment groups) was rejected at \**p* < 0.05, \*\**p* < 0.01, \*\*\**p* < 0.001 or \*\*\*\**p* < 0.0001. The error bars represent standard deviation (n=7).

Vaccination with Hsp18 recombinant protein alone or Hsp18 + R_4_Pam_2_Cys induced Hsp18-specific antibody titres in both strains of mice (Fig. 3C, D). Vaccine boost with Hsp18 recombinant protein alone induced significantly higher Hsp18-specific antibody responses in BALB/c mice compared to a single vaccination with Hsp18 protein (*p* < 0.0001). Boosting with protein alone in C57BL/6 did not significantly increase antibody titres. Hsp18 + R_4_Pam_2_Cys induced Hsp18-specific antibody responses in both mouse strains after primary vaccination (*p* < 0.0001 in BALB/c and *p* = 0.0165 in C57BL/6) and the Hsp18-specific antibody titre significantly increased after booster vaccination (*p* < 0.0001 in BALB/c and p = 0.0016 in C57BL/6). In all strains, the antibody titres induced by Hsp18 + R_4_Pam_2_Cys were significantly higher than vaccination with Hsp18 protein alone (*p* = 0.0004 in BALB/c and *p* < 0.0001 in C57BL/6) (Fig. 3C, D) with negligible levels of antibodies seen in mice vaccinated with only R_4_Pam_2_Cys, or BCG.

### Measurement of IgG antibody subtypes following MUL_3720 + R_4_Pam_2_Cys and Hsp18 + R_4_Pam_2_Cys vaccination

Quantifying levels of IgG antibody shows that the predominant isotypes produced by MUL_3720 were IgG_1_ and IgG2_b_ (Fig. 3E) with no significant difference between these isotype titres. Vaccination with MUL_3720 + R_4_Pam_2_Cys produces significantly more IgG_1_ and IgG2_b_ antibodies (*p =* 0.0076). The antibody titres for both isotypes were highest prior to infection with *M. ulcerans* (day 33) and decreased after infection by day 63. This vaccine was capable of inducing IgG2a antibodies, which was detected also on day 33, however in smaller amounts than IgG_1_ and IgG2_b_ (*p* = 0.0399 for MUL_3720 + R_4_Pam_2_Cys) (Fig. 3E).

Similar to vaccination with MUL_3720, Hsp18 was also capable of inducing strong IgG antibody titres. The predominant isotype was IgG_1_ which Hsp18 + R_4_Pam_2_Cys elicited more than any other isotype (Fig. 3F) including IgG2_a_ and IgG2_b_, Again, these titres was highest at day 33 and decreased significantly after infection on day 63. This trend was also observed after vaccination with Hsp18 alone (*p* = 0.0018 vs IgG2_a_ and *p =* 0.0076 vs IgG2_b_, respectively at day 33).

### MUL_3720 + R_4_Pam_2_Cys and Hsp18 + R_4_Pam_2_Cys do not protect against the onset of BU

As both vaccines were capable of inducing protein-specific antibody responses, they were tested in a murine challenge model to measure their protective efficacy. Efficacy was measured by time delay to the onset of ulceration in a mouse tail infection model. There are a progression of clinical symptoms for Buruli ulcer in this model (Fig. 4). Once ulceration has been reached the disease would likely continue until the tail became necrotic. Therefore, the experimental endpoint was deemed to be the point of ulceration.

**Figure 4.**
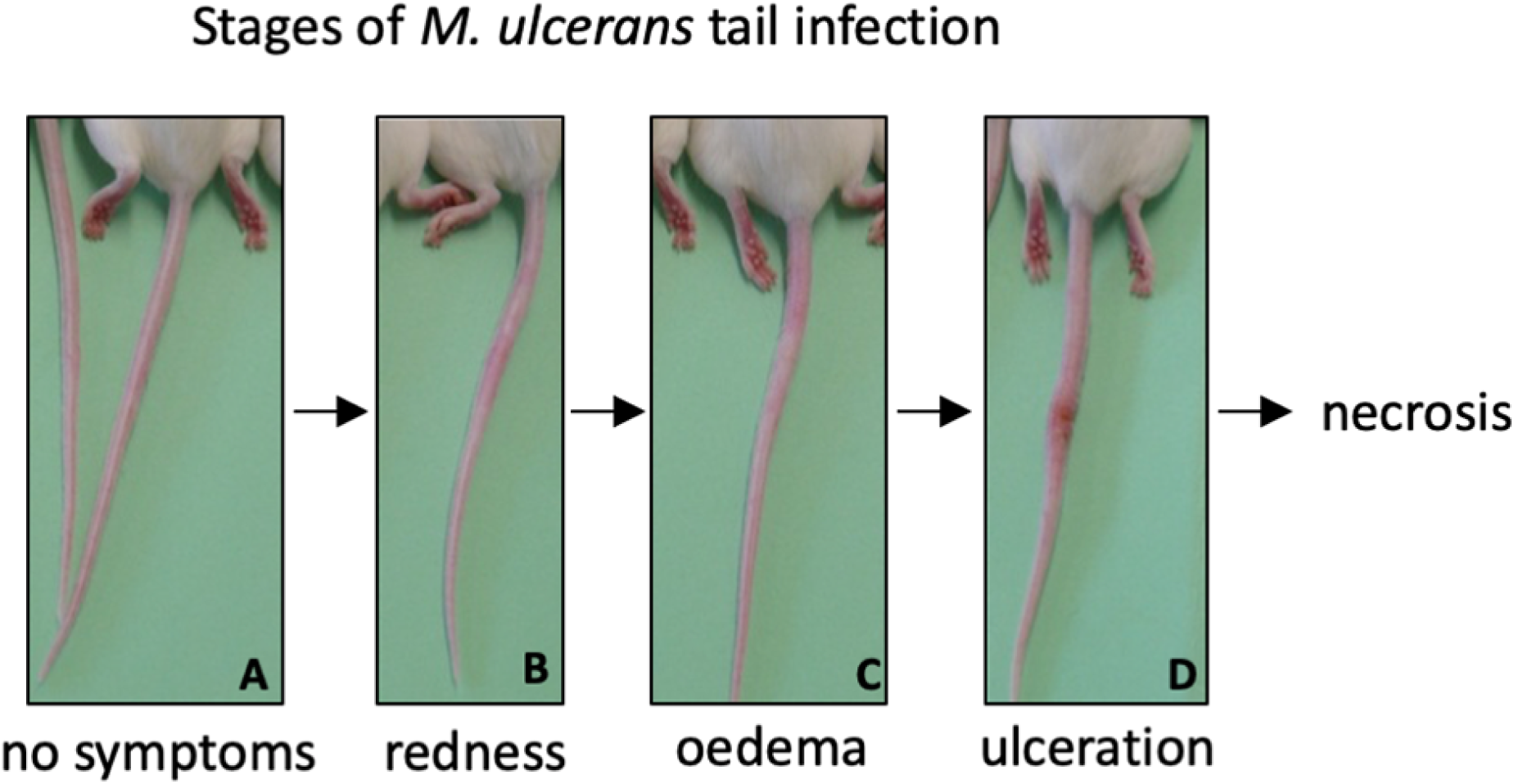
Progression of BU in the murine tail infection model over time. **(A)** Healthy mouse tail. (**B)** Appearance of a small sign of redness at the site of tail infection. **(C)** Oedema surrounding the initial site of redness. **(D)** Tail lesion at the point of ulceration. This is typically identified by excessive oedema and redness at the site of imminent ulceration. Mice were culled before ulcerative lesions appeared.

After the scheduled vaccinations, mice were challenged via subcutaneous tail inoculation with 1 × 10^4^ CFU of *M. ulcerans* and observed for up to 40 days. In BALB/c and C57BL/6 mice there was no significant difference between the time to ulceration between control mice (mice not vaccinated with recombinant protein, such as R_4_Pam_2_Cys alone and BCG) and mice vaccinated with either MUL_3720 + R_4_Pam_2_Cys or Hsp18 + R_4_Pam_2_Cys (Figure 5A and 5B). There was also no significant difference in the time to ulceration between mice that were vaccinated with MUL_3720 + R_4_Pam_2_Cys or Hsp18 + R_4_Pam_2_Cys and BCG, the benchmark for mycobacterial vaccine efficacy. Signs of infection in all BALB/c and C57BL/6 mice were visible by day 33 (Table 1 and Table 2) and all mice reached ulceration by day 63, 30 days post-*M. ulcerans* challenge (Fig. 5A, B).

**Figure 5.**
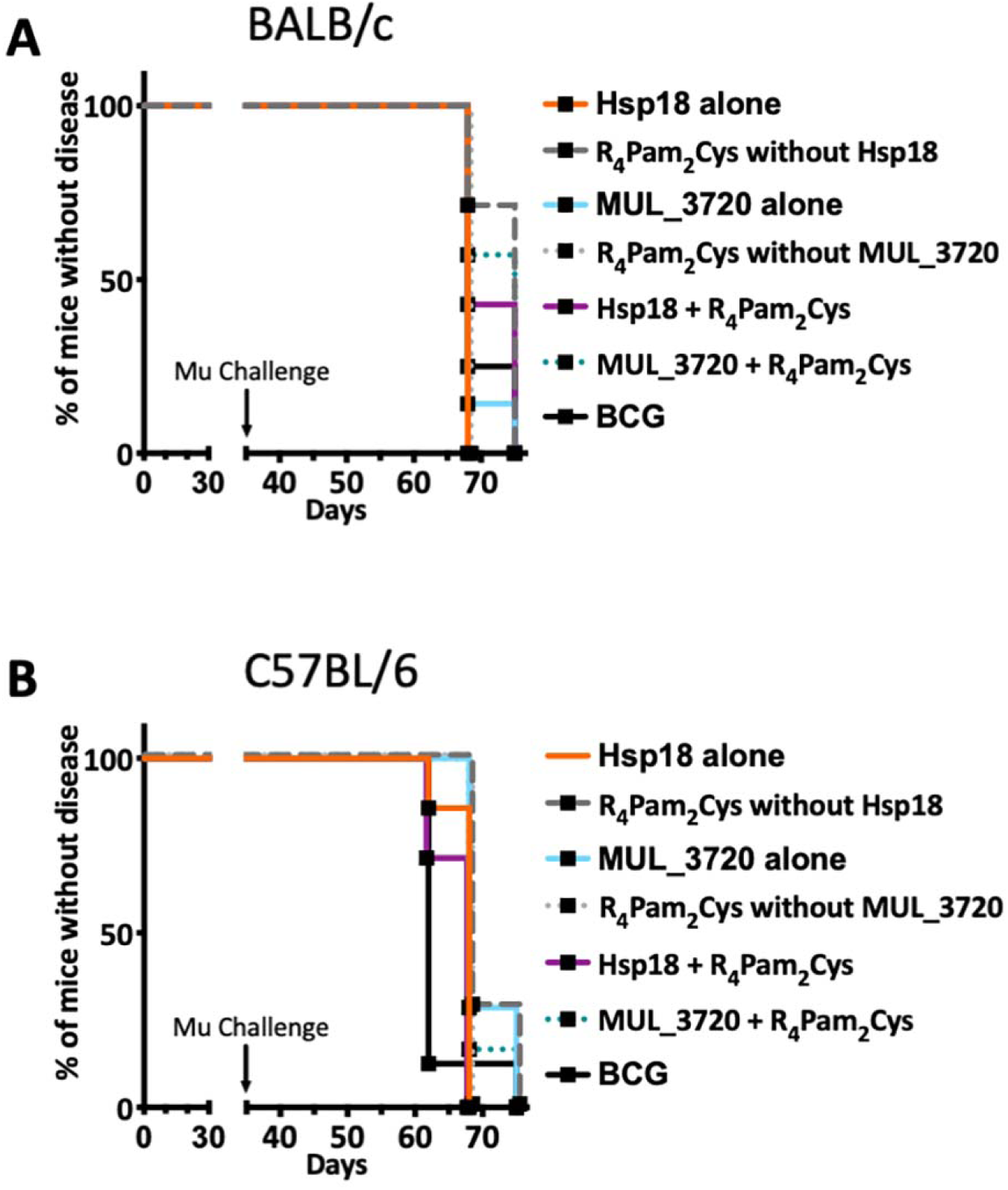
Vaccine performance using murine tail infection model of BU. Survival analysis showing the time taken (days) for each mouse to reach ulceration for different vaccination groups post *M. ulcerans* challenge. **(A)** BALB/c mice (n=7) and (**B)** C57BL/6 mice (n=7). The null hypothesis (no difference in mean antibody responses between treatment groups) was rejected if \**p* < 0.05, \*\**p* < 0.01, \*\*\**p* < 0.001 or \*\*\*\**p* < 0.0001.

### Antibody titres do not correlate with protection against *M. ulcerans*

High antibody titres were observed in all mice vaccinated with either recombinant MUL_3720 or Hsp18, particularly in the secondary response after booster vaccination (Fig. 3A-D) prior to *M. ulcerans* challenge. However, mice vaccinated with protein alone or protein plus lipopeptide adjuvant all succumbed to infection by day 75. The sera from mice at the day 63 was used to quantify antibody titres during infection. At day 63 all mice still had detectable protein-specific antibodies against the recombinant protein with which they were vaccinated (Figure 3A-3D). In BALB/c mice (Fig. 3A, C) the antibody titres at day 63 were lower than after the secondary response prior to challenge (*p* < 0.0001 for both Hsp18 + R_4_Pam_2_Cys and MUL_3720 + R_4_Pam_2_Cys) but remained significantly higher than at day 0 (*p* < 0.0001 for both Hsp18 + R_4_Pam_2_Cys and MUL_3720 + R_4_Pam_2_Cys). In C57BL/6 mice (Fig. 3B and 3D), antibody titres against MUL_3720 or Hsp18 from mice vaccinated with either protein alone or protein plus lipopeptide adjuvant were also significantly decreased at day 63 compared to the secondary response at day 35 (*p* < 0.0001 and *p* = 0.0406 for MUL_3720 + R_4_Pam_2_Cys and Hsp18 + R_4_Pam_2_Cys, respectively). Similar to BALB/c mice, the day 63 respective protein-specific antibodies for MUL_3720 + R_4_Pam_2_Cys and Hsp18 + R_4_Pam_2_Cys were significantly higher than at day 0 (*p* < 0.0001 and *p* = 0.0004 for MUL_3720 + R_4_Pam_2_Cys and Hsp18 + R_4_Pam_2_Cys, respectively).

### Challenge with *M. ulcerans* did not induce protein-specific antibody levels comparable to vaccination with MUL_3720 or Hsp18

MUL_3720 and Hsp18 recombinant proteins are immunogenic and capable of inducing protein-specific antibody responses after vaccination. However, only minor detectable antibody responses against either recombinant MUL_3720 or Hsp18 at day 63 (Fig. 3A-D) were found in micr vaccinated with R_4_Pam_2_Cys alone or BCG then challenged with *M. ulcerans*. These responses are much lower than the protein-specific antibody responses generated from MUL_3720 or Hsp18 vaccinated mice, particularly in C57/BL6 mice (*p* < 0.0001) (Fig. 6). Animals from both mouse strains that were vaccinated with R_4_Pam_2_Cys alone or BCG showed no increase in protein-specific antibody responses against either recombinant MUL_3720 and Hsp18 on day 63 post-*M. ulcerans* challenge (Fig. 3A-D), even though these two proteins are both expressed in *M. ulcerans*.

**Figure 6.**
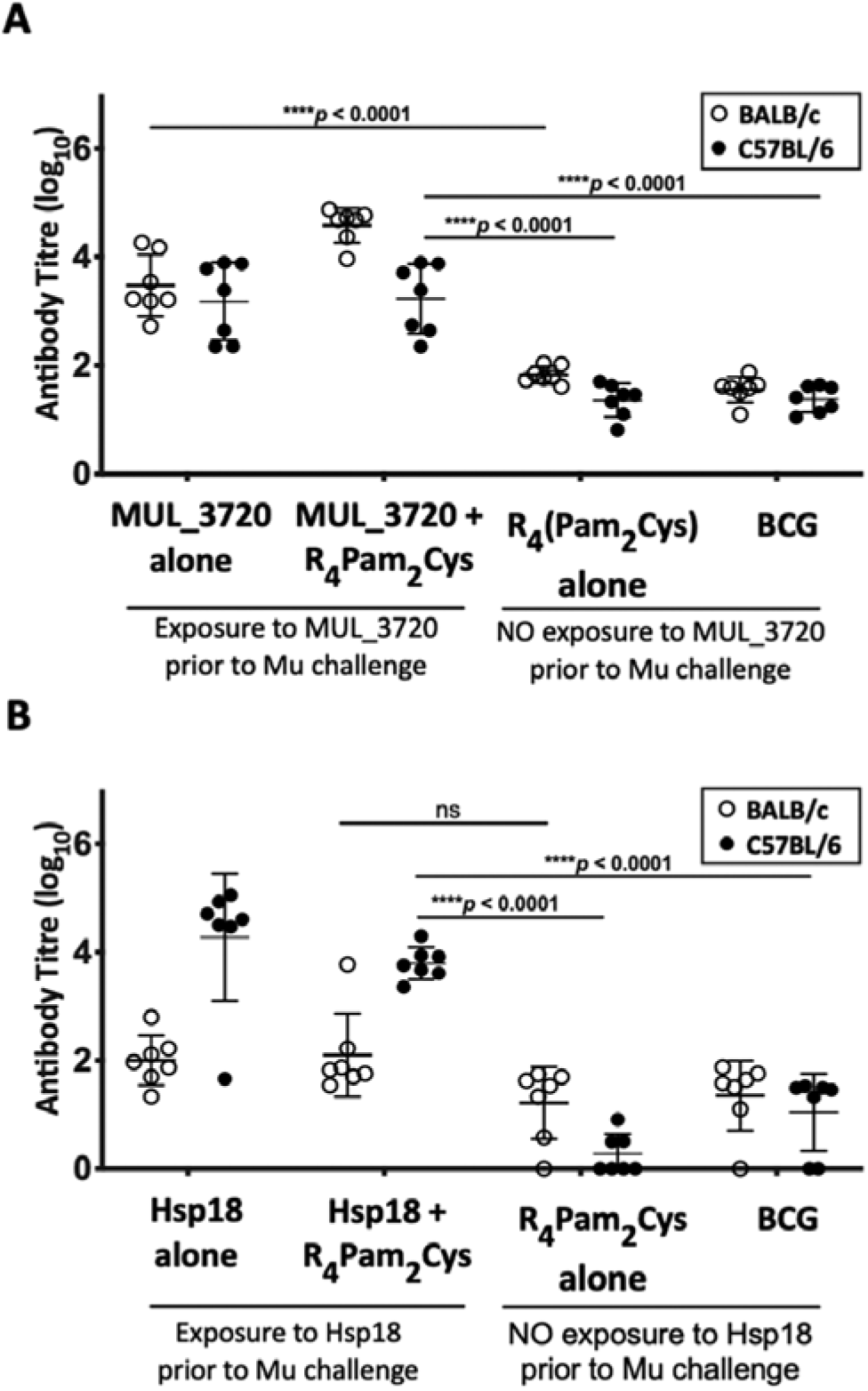
Antibody responses to recombinant MUL_3720 and Hsp18 in unvaccinated mice. **(A)** MUL_3720-specific antibody titres and **(B)** Hsp18-specific antibody titres from BALB/c mice (clear circles) and C57BL/6 mice (black circles) at day 63 (post-MU exposure). The null hypothesis (no difference in mean antibody responses between treatment groups) was rejected if \**p* <0.05, \*\**p* <0.01, \*\*\**p* <0.001 or \*\*\*\**p* <0.0001. The error bars are standard deviation (n=7).

## Discussion

This study aimed to develop a vaccine against *M. ulcerans* utilizing two previously described cell-wall associated proteins, Hsp18 and MUL_3720 [40-43]. Both the MUL_3720 and Hsp18-based vaccines were capable of inducing high antibody titres, but these responses were not associated with protection (Fig. 5). This may indicate that these proteins, while strongly immunogenic, play no major role in pathogenesis, so targeting them with potentially neutralizing antibodies induced by the vaccine has no impact on disease. Alternatively, antibodies raised by these vaccines may not have had the functional potential to control infection. In addition to antigen binding, antibodies engage via their Fc domains with Fcγ receptors (FcγR) present on innate immune cells (NK cells, monocytes, macrophages and neutrophils) to rapidly recruit the anti-microbial activity of the innate immune system. Antibodies with these functions can promote control of a pathogen through the activation of multiple effector cell functions, including Ab dependent cellular cytotoxicity, cellular phagocytosis and/or cytokine and enzyme secretion [55-57]. Recent research has shown that mice lacking antibodies have increased susceptibility to *M. tuberculosis* infection [58] and non-human primates treated to deplete B cells also exhibit increased bacterial burden [59]. Despite the findings of this research, it is likely that B cells and antibody responses still play a role in controlling *M. ulcerans* infection in this model, albeit with different specificities. Future research could use human BU patient cohorts and mouse infection models to attempt to characterize the targets, functional and structural aspects of antibody responses that differentiate subjects able to control BU from susceptible subjects. It might then be possible to use B cell probe technologies to isolate Ag-specific memory B cells from individuals that control *M. ulcerans* infection and then clone the immunoglobulin gene sequences identified [60]. Antigen-specific monoclonal Abs (mAbs) could then be generated and characterized for their *in vitro* anti-microbial activity and used in *in vivo* mouse passive transfer studies to determine potential use as mAb therapeutics against BU.

Another explanation for the ineffectiveness of antibodies in this study may be due to the localized immune suppression induced by the *M. ulcerans* toxin mycolactone at the site of infection. Mycolactone diffuses into tissue surrounding the bacteria [61-63]. Mycolactone is a cytotoxin that modulates the function of several immune cells [63, 64]. The toxin inhibits the Sec61 translocon, affecting T cell activation, impairing T cell responsiveness and distorting cytokine production [62, 63]. The mycolactone-induced depletion of T cell homing to peripheral lymph nodes affects subsequent B-cell activation and migration from the lymphatics [65]. The antibodies induced by the vaccine in this study may be functional but unable to access bacteria within the infection or it may be that multiple effector cell functions have been modulated by mycolactone exposure through interference with receptor expression on key innate immune cells, rendering these cells poorly responsive to antibodies. Suppression of protein-specific antibody production in the presence of mycolactone has been observed [66]. Mycolactone administered to a different location to the antigen caused no reduction to systemic antigen-specific IgG titres [66], similar to the observations from our study.

The greatest antibody responses were of the IgG1 subclass. Typical antibody responses against proteins occur via B cell isotype switching from IgM (non-specific antibody isotype) to IgG. There are 4 subclasses of IgG (IgG1, IgG2, IgG3 and IgG4) and isotype switching to predominantly IgG1 suggests refinement of immune responses to respond specifically to either MUL_3720 or Hsp18, as IgG1 is capable of binding to protein antigens [67]. IgG1 can also bind all forms of FcγR which is required to elicit and mediate effector immune functions as described above [68]. The presence of IgG2 suggest further isotype switching from IgG1 to IgG2_a/b_ as the immune response develops. IgG2 is less effective at inducing phagocytosis and fixing complement and is more commonly associated with polysaccharide antigens. Though tests on the recombinant proteins had undetectable levels of lipopolysaccharide (LPS), there could be trace amounts from the *E. coli* expression vector boosting IgG2 responses. Studies analysing antibodies generated during leprosy and TB infection show a switch from IgG1 to IgG2 antibodies for leprosy and a persistence of IgG1 and IgG3 antibodies for TB [69]. As isotype switching of antibodies requires help by T helper cells, future work could therefore also incorporate studies on the effect of vaccination and subsequent *M. ulcerans*-infection on T cells as well as antibody responses.

In this study, all mice succumbed to infection in a relatively short period (40 days) compared to previous mouse tail infection models [70] and human BU, where the incubation period is estimated at 4.8 months before the onset of ulceration [71]. All BALB/c and C57BL/6 mice succumbed to infection by 40 days after MU infection, even mice that were vaccinated by *M. bovis* BCG. *M. bovis* BCG has been previously shown to delay the onset of disease on average by at least 6 weeks [21, 22, 31]. In this study however, there was no significant difference between mice vaccinated with either MUL_3720 or Hsp18 protein alone or with both proteins plus R_4_Pam_2_Cys. This suggests that *M. bovis* BCG is ineffective at protecting mice in this model of *M. ulcerans* vaccination. This failure to observe any protective impact of *M. bovis* BCG might be a reflection of the challenge strain of *M. ulcerans* used (strain Mu_1G897) and/or the high challenge dose used (10^4^ bacteria). High concentrations (>10^4^ bacteria) have not been reported in environmental sources of *M. ulcerans* [9, 72-75], consistent with the hypothesis that a relatively small bacterial inoculum is required to establish BU [9]. At the time this study was conducted the minimum infectious dose (ID_50_) for BU had not been determined, however the ID_50_ has since been identified as approximately 3 CFU [48]. Future studies should therefore use a murine model that is more representative of a natural *M. ulcerans* infection, reflected both in the mode of *M. ulcerans* entry into the subcutaneous tissue and in the dose of bacteria used for challenge.

## Conclusions

Vaccination with either MUL_3720 or Hsp18 proteins induced high antibody titres. These responses were augmented when either protein was linked with the lipopeptide adjuvant R_4_Pam_2_Cys. However, robust antibody responses did not correlate with protection against challenge with *M. ulcerans*. Future work could test different *M. ulcerans* antigens in vaccine formulations against Buruli ulcer. As mycolactone is a key virulence factor, neutralising this toxin early in infection by targeting the PKS enzymes required for its biosynthesis could be a focus for future vaccination developments. Using a low *M. ulcerans* inoculum as a more realistic vaccine challenge dose is also warranted.

## Acknowledgements

We thank Roy Robins-Browne for providing the pET-30b MOD plasmid used for the expression of recombinant Hsp18.

## References

1. Guarner, J., et al., Histopathologic features of Mycobacterium ulcerans infection. Emerg Infect Dis, 2003. 9(6): p. 651–656.

2. Vincent, Q.B., et al., Clinical epidemiology of laboratory-confirmed Buruli ulcer in Benin: a cohort study. Lancet Glob Health, 2014. 2(7): p. e422–30.

3. Hayman, J. and A. McQueen, The pathology of Mycobacterium ulcerans infection. Pathology, 1985. 17(4): p. 594–600.

4. Oliveira, M.S., et al., Infection with Mycobacterium ulcerans induces persistent inflammatory responses in mice. (0019-9567 (Print)).

5. Woodring, J.H., et al., Update: the radiographic features of pulmonary tuberculosis. American Journal of Roentgenology, 1986. 146(3): p. 497–506.

6. van der Werf, T.S., et al., Mycobacterium ulcerans infection. Lancet, 1999. 354(9183): p. 1013–8.

7. Marsollier, L., et al., Aquatic insects as a vector for Mycobacterium ulcerans. Appl Environ Microbiol, 2002. 68(9): p. 4623–8.

8. Meyers, W.M., et al., Human Mycobacterium ulcerans infections developing at sites of trauma to skin. Am J Trop Med Hyg, 1974. 23(5): p. 919–23.

9. Stinear, T., et al., Identification of Mycobacterium ulcerans in the environment from regions in Southeast Australia in which it is endemic with sequence capture-PCR. Appl Environ Microbiol, 2000. 66(8): p. 3206–13.

10. Michael, J.L., et al., Epidemiology of Buruli Ulcer Infections, Victoria, Australia, 2011– 2016. Emerging Infectious Disease journal, 2018. 24(11): p. 1988.

11. Organization, W.H. Buruli ulcer - number of new reported cases. Global Health Observatory data repository 2019 19 June 2019 [cited 2020 29 January]; Available from: http://apps.who.int/gho/data/node.main.A1631.

12. Simpson, H., et al., Mapping the global distribution of Buruli ulcer: a systematic review with evidence consensus. The Lancet Global Health, 2019. 7(7): p. e912–e922.

13. Omansen, T.F., et al., Global Epidemiology of Buruli Ulcer, 2010–2017, and Analysis of 2014 WHO Programmatic Targets. Emerging Infectious Diseases, 2019. 25(12): p. 2183–2190.

14. Yerramilli, A., et al., The location of Australian Buruli ulcer lesions—Implications for unravelling disease transmission. PLOS Neglected Tropical Diseases, 2017. 11(8): p. e0005800.

15. Sarfo, F.S., et al., Clinical efficacy of combination of rifampin and streptomycin for treatment of Mycobacterium ulcerans disease. Antimicrob Agents Chemother, 2010. 54(9): p. 3678–85.

16. van der Werf, T.S., et al., Mycobacterium ulcerans disease. Bull World Health Organ, 2005. 83(10): p. 785–91.

17. Guarner, J., Buruli Ulcer: Review of a Neglected Skin Mycobacterial Disease. Journal of clinical microbiology, 2018. 56(4): p. e01507–17.

18. Herbinger, K.H., et al., Excision of Pre-Ulcerative Forms of Buruli Ulcer Disease: A Curative Treatment? Infection, 2008.

19. Boyd, S.C., et al., Epidemiology, clinical features and diagnosis of Mycobacterium ulcerans in an Australian population. Med J Aust, 2012. 196(5): p. 341–4.

20. Amofah, G., et al., Buruli ulcer in Ghana: results of a national case search. Emerg Infect Dis, 2002. 8(2): p. 167–70.

21. Tanghe, A., et al., Protective Efficacy of a DNA Vaccine Encoding Antigen 85A from Mycobacterium bovis BCG against Buruli Ulcer. Infect Immun, 2001. 69(9): p. 5403–5411.

22. Tanghe, A., et al., A booster vaccination with Mycobacterium bovis BCG does not increase the protective effect of the vaccine against experimental Mycobacterium ulcerans infection in mice. Infect Immun, 2007. 75(5): p. 2642–4.

23. Phillips, R.O., et al., Effectiveness of routine BCG vaccination on buruli ulcer disease: a case-control study in the Democratic Republic of Congo, Ghana and Togo. PLoS Negl Trop Dis, 2015. 9(1): p. e3457.

24. Group, T.U.B., BCG vaccination against mycobacterium ulcerans infection (Buruli ulcer). First results of a trial in Uganda. Lancet, 1969. 293(7586): p. 111–5.

25. Smith, P.G., et al., The protective effect of BCG against Mycobacterium ulcerans disease: a controlled trial in an endemic area of Uganda. Trans R Soc Trop Med Hyg, 1976. 70(5-6): p. 449–57.

26. Tanghe, A., et al., Improved protective efficacy of a species-specific DNA vaccine encoding mycolyl-transferase Ag85A from Mycobacterium ulcerans by homologous protein boosting. PLoS Negl Trop Dis, 2008. 2(3): p. e199.

27. Coutanceau, E., et al., Immunogenicity of Mycobacterium ulcerans Hsp65 and protective efficacy of a Mycobacterium leprae Hsp65-based DNA vaccine against Buruli ulcer. Microbes Infect, 2006. 8(8): p. 2075–81.

28. Roupie, V., et al., Analysis of the vaccine potential of plasmid DNA encoding nine mycolactone polyketide synthase domains in Mycobacterium ulcerans infected mice. PLoS Negl Trop Dis, 2014. 8(1): p. e2604.

29. Bolz, M., et al., Use of Recombinant Virus Replicon Particles for Vaccination against Mycobacterium ulcerans Disease. PLoS Negl Trop Dis, 2015. 9(8): p. e0004011.

30. Bolz, M., et al., Vaccination with the Surface Proteins MUL_2232 and MUL_3720 of Mycobacterium ulcerans Induces Antibodies but Fails to Provide Protection against Buruli Ulcer. PLoS Negl Trop Dis, 2016. 10(2): p. e0004431.

31. Fraga, A.G., et al., Cellular immunity confers transient protection in experimental Buruli ulcer following BCG or mycolactone-negative Mycobacterium ulcerans vaccination. PLoS One, 2012. 7(3): p. e33406.

32. Watanabe, M., et al., Protective effect of a dewaxed whole-cell vaccine against Mycobacterium ulcerans infection in mice. Vaccine, 2015. 33(19): p. 2232–9.

33. Hart, B.E., L.P. Hale, and S. Lee, Immunogenicity and protection conferred by a recombinant Mycobacterium marinum vaccine against Buruli ulcer. Trials in Vaccinology, 2016. 5: p. 88–91.

34. Converse, P.J., et al., BCG-mediated protection against Mycobacterium ulcerans infection in the mouse. PLoS Negl Trop Dis, 2011. 5(3): p. e985.

35. Hart, B.E., L.P. Hale, and S. Lee, Recombinant BCG Expressing Mycobacterium ulcerans Ag85A Imparts Enhanced Protection against Experimental Buruli ulcer. PLoS Negl Trop Dis, 2015. 9(9): p. e0004046.

36. Hart, B.E. and S. Lee, Overexpression of a Mycobacterium ulcerans Ag85B-EsxH Fusion Protein in Recombinant BCG Improves Experimental Buruli Ulcer Vaccine Efficacy. PLoS Negl Trop Dis, 2016. 10(12): p. e0005229.

37. Trigo, G., et al., Phage therapy is effective against infection by Mycobacterium ulcerans in a murine footpad model. PLoS Negl Trop Dis, 2013. 7(4): p. e2183.

38. Siegrist, C.-A., 2 - Vaccine Immunology, in Plotkin’s Vaccines (Seventh Edition), S.A. Plotkin, et al., Editors. 2018, Elsevier. p. 16-34.e7.

39. Plotkin, S.A., Vaccination against the major infectious diseases. Comptes Rendus de l’Académie des Sciences - Series III - Sciences de la Vie, 1999. 322(11): p. 943–951.

40. Pidot, S.J., et al., Serological evaluation of Mycobacterium ulcerans antigens identified by comparative genomics. PLoS Negl Trop Dis, 2010. 4(11): p. e872.

41. Pidot, S.J., et al., Regulation of the 18 kDa heat shock protein in Mycobacterium ulcerans: an alpha-crystallin orthologue that promotes biofilm formation. Mol Microbiol, 2010. 78(5): p. 1216–31.

42. Vettiger A, S.N., Ruf M-T, Röltgen K, Pluschke G., Localization of Mycobacterial Antigens by Immunofluorescence Staining of Agarose Embedded Cells. J Mycobact Dis., 2014. 4(3): p. 150.

43. Dreyer, A., et al., Identification of the Mycobacterium ulcerans protein MUL_3720 as a promising target for the development of a diagnostic test for Buruli ulcer. PLoS neglected tropical diseases, 2015. 9(2): p. e0003477–e0003477.

44. Chua, B.Y., et al., The use of a TLR2 agonist-based adjuvant for enhancing effector and memory CD8 T-cell responses. Immunol Cell Biol, 2014. 92(4): p. 377–83.

45. Chua, B.Y., et al., Soluble proteins induce strong CD8+ T cell and antibody responses through electrostatic association with simple cationic or anionic lipopeptides that target TLR2. J Immunol, 2011. 187(4): p. 1692–701.

46. Chua, B.Y., et al., Hepatitis C VLPs Delivered to Dendritic Cells by a TLR2 Targeting Lipopeptide Results in Enhanced Antibody and Cell-Mediated Responses. PLOS ONE, 2012. 7(10): p. e47492.

47. Christiansen, D., et al., Antibody Responses to a Quadrivalent Hepatitis C Viral-Like Particle Vaccine Adjuvanted with Toll-Like Receptor 2 Agonists. Viral Immunology, 2018. 31(4): p. 338–343.

48. Wallace, J.R., et al., Mycobacterium ulcerans low infectious dose and mechanical transmission support insect bites and puncturing injuries in the spread of Buruli ulcer. PLoS Negl Trop Dis, 2017. 11(4): p. e0005553.

49. Laemmli, U.K., Cleavage of structural proteins during the assembly of the head of bacteriophage T4. Nature, 1970. 227(5259): p. 680–5.

50. Sekiya, T., et al., PEGylation of a TLR2-agonist-based vaccine delivery system improves antigen trafficking and the magnitude of ensuing antibody and CD8(+) T cell responses. Biomaterials, 2017. 137 (1878-5905 (Electronic)): p. 61–72.

51. Wijayadikusumah, A.R., et al., Structure-function relationships of protein-lipopeptide complexes and influence on immunogenicity. Amino Acids, 2017. 49(10): p. 1691–1704.

52. Wijayadikusumah, A.R., An evaluation of charged Pam2Cys-based lipopeptides as novel adjuvants for subunit-based vaccines. 2017.

53. Tan, A.C., et al., Intranasal Administration of the TLR2 Agonist Pam2Cys Provides Rapid Protection against Influenza in Mice. Mol Pharm, 2012.

54. Mifsud, E.J., et al., Reducing the impact of influenza-associated secondary pneumococcal infections. Immunol Cell Biol, 2016. 94(1): p. 101–8.

55. Lu, L.L., et al., A Functional Role for Antibodies in Tuberculosis. Cell, 2016. 167(2): p. 433–443 e14.

56. Damelang, T., et al., Role of IgG3 in Infectious Diseases. Trends Immunol, 2019. 40(3): p. 197–211.

57. Chung, A.W., et al., Dissecting Polyclonal Vaccine-Induced Humoral Immunity against HIV Using Systems Serology. Cell, 2015. 163(4): p. 988–98.

58. Maglione, P.J., J. Xu, and J. Chan, B cells moderate inflammatory progression and enhance bacterial containment upon pulmonary challenge with Mycobacterium tuberculosis. J Immunol, 2007. 178(11): p. 7222–34.

59. Phuah, J., et al., Effects of B Cell Depletion on Early Mycobacterium tuberculosis Infection in Cynomolgus Macaques. Infect Immun, 2016. 84(5): p. 1301–1311.

60. McLean, M.R., et al., Dimeric Fcgamma Receptor Enzyme-Linked Immunosorbent Assay To Study HIV-Specific Antibodies: A New Look into Breadth of Fcgamma Receptor Antibodies Induced by the RV144 Vaccine Trial. J Immunol, 2017. 199(2): p. 816–826.

61. George, K.M., et al., Mycolactone: a polyketide toxin from Mycobacterium ulcerans required for virulence. Science, 1999. 283(5403): p. 854–7.

62. Boulkroun, S., et al., Mycolactone suppresses T cell responsiveness by altering both early signaling and posttranslational events. J Immunol, 2010. 184(3): p. 1436–44.

63. Baron, L., et al., Mycolactone subverts immunity by selectively blocking the Sec61 translocon. J Exp Med, 2016. 213(13): p. 2885–2896.

64. Ogbechi, J., et al., Inhibition of Sec61-dependent translocation by mycolactone uncouples the integrated stress response from ER stress, driving cytotoxicity via translational activation of ATF4. Cell Death & Disease, 2018. 9(3): p. 397.

65. Guenin-Mace, L., et al., Mycolactone impairs T cell homing by suppressing microRNA control of L-selectin expression. Proc Natl Acad Sci U S A, 2011. 108(31): p. 12833–8.

66. Shinoda, N., H. Nakamura, and M. Watanabe, Suppressive effect of mycolactone-containing fraction from Mycobacterium ulcerans on antibody production against coadministered antigens. Biomedical Research and Clinical Practice, 2017. 2(1): p. 1–6.

67. Schroeder, H.W., Jr. and L. Cavacini, Structure and function of immunoglobulins. The Journal of allergy and clinical immunology, 2010. 125(2 Suppl 2): p. S41–S52.

68. Sibéril, S., et al., FcγR: The key to optimize therapeutic antibodies? Critical Reviews in Oncology/Hematology, 2007. 62(1): p. 26–33.

69. Sousa, A.O., et al., IgG subclass distribution of antibody responses to protein and polysaccharide mycobacterial antigens in leprosy and tuberculosis patients. Clinical and experimental immunology, 1998. 111(1): p. 48–55.

70. Omansen, T.F., et al., In Vivo Imaging of Bioluminescent Mycobacterium ulcerans: A Tool to Refine the Murine Buruli Ulcer Tail Model. The American Journal of Tropical Medicine and Hygiene, 2019. 101(6): p. 1312–1321.

71. Loftus, M.J., et al., The incubation period of Buruli ulcer (Mycobacterium ulcerans infection) in Victoria, Australia - Remains similar despite changing geographic distribution of disease. PLoS Negl Trop Dis, 2018. 12(3): p. e0006323.

72. Fyfe, J.A., et al., A major role for mammals in the ecology of Mycobacterium ulcerans. PLoS Negl Trop Dis, 2010. 4(8): p. e791.

73. Williamson, H.R., et al., Detection of Mycobacterium ulcerans in the Environment Predicts Prevalence of Buruli Ulcer in Benin. PLOS Neglected Tropical Diseases, 2012. 6(1): p. e1506.

74. Marion, E., et al., Seasonal and regional dynamics of M. ulcerans transmission in environmental context: deciphering the role of water bugs as hosts and vectors. PLoS neglected tropical diseases, 2010. 4(7): p. e731–e731.

75. Johnson, P.D., et al., Mycobacterium ulcerans in mosquitoes captured during outbreak of Buruli ulcer, southeastern Australia. Emerg Infect Dis, 2007. 13(11): p. 1653–60.

