## Supplementary material for "High antibody titres induced by protein subunit vaccines against Buruli ulcer using *Mycobacterium ulcerans* antigens Hsp18 and MUL_3720": Table S1

**Supplementary Table 1. Summary of protein antigen and assay characteristics**

| **Protein** | **Size** | **Function** | **Expression**  **Vector** | **Presence of histidine tag (HIS-tag)** | **Endotoxin concentration** | **Putative charge^a^** | **Preferentially bound lipopeptide adjuvant** |
| --- | --- | --- | --- | --- | --- | --- | --- |
| MUL_3720 | 22kDa | Putative cell-wall associated protein | pDest17 | N-terminal 6x histidine tag | Below detectable limit | -13.7 | R_4_Pam_2_Cys at 1:5 protein to lipopeptide ratio |
| Hsp18 | 18kDa | Heat shock protein | pET-30b MOD^b^ | N-terminal 6x histidine tag | Below detectable limit | -5.8 | R_4_Pam_2_Cys at 1:5 protein to lipopeptide ratio |

^a^ As calculated by <https://pepcalc.com/protein-calculator.php> (Innovagen) using translated amino acid sequence from DNA sequence.

^b^ Commercial Novagen pET-30b vector was modified to exclude DNA sequence from the end of the *Nde*I restriction site and three bases from the beginning of the *Eco*RV restriction site.
